# A novel hamster model of SARS-CoV-2 respiratory infection using a pseudotyped virus

**DOI:** 10.1101/2021.09.17.460745

**Authors:** Hiroshi Yamada, Soichiro Sasaki, Hideki Tani, Mayu Somekawa, Hitoshi Kawasuji, Yumiko Saga, Yoshihiro Yoshida, Yoshihiro Yamamoto, Yoshihiro Hayakawa, Yoshitomo Morinaga

**Author notes:** **Corresponding author** Yoshitomo Morinaga, MD, PhD, Department of Microbiology, Graduate School of Medicine and Pharmaceutical Sciences, University of Toyama, 2630 Sugitani, Toyama, 930-0194 Japan.

## Abstract

**Background:** Severe acute respiratory syndrome coronavirus 2 (SARS-CoV-2) is a biosafety level (BSL)-3 pathogen; therefore, its research environment is strictly limited. Pseudotyped viruses that mimic SARS-CoV-2 have been widely used for *in vitro* evaluation because they are available in BSL-2 containment laboratories; however, *in vivo* application is inadequate. Therefore, animal models that can be instigated with animal BSL-2 will increase opportunities for *in vivo* evaluations.

**Methods:** Hamsters (6-to 10-week-old males) were intratracheally inoculated with luciferase-expressing vesicular stomatitis virus (VSV)-based SARS-CoV-2 pseudotyped virus. The lungs were harvested 24 h after inoculation, and luminescence was measured using an *in vivo* imaging system.

**Results:** Lung luminescence after inoculation with the SARS-CoV-2 pseudotyped virus increased in a dose-dependent manner. VSV-G (envelope [G]) pseudotyped virus also induced luminescence; however, a 100-fold concentration was required to reach a level similar to that of the SARS-CoV-2 pseudotyped virus.

**Conclusions:** The SARS-CoV-2 pseudotyped virus is applicable to SARS-CoV-2 respiratory infections in a hamster model. Because of the single-round infectious virus, the model can be used to study the steps from viral binding to entry, which will be useful for future research regarding SARS-CoV-2 entry without using live SARS-CoV-2 or transgenic animals.

## Introduction

Emerging severe acute respiratory syndrome coronavirus 2 (SARS-CoV-2) causes coronavirus disease 2019 (COVID-19) and has seriously affected our health and social activities. There is an urgent need to elucidate the pathophysiology of COVID-19 and develop treatment and prophylactic approaches.

SARS-CoV-2 is classified as a biosafety level (BSL)-3 pathogen and requires a limited laboratory environment and strict control by skilled researchers [1]. Research using live viruses has the advantage of evaluating disease pathogenicity more directly; however, there are some limitations regarding laboratories and effort. Therefore, pseudotyped viruses, which are single-round infectious virus particles with an envelope protein originating from a different virus [2], have been widely used in COVID-19-related research because they can be used in BSL-2 containment laboratories. As an alternative to live SARS-CoV-2, we previously developed a pseudotyped virus, vesicular stomatitis virus (VSV) expressing luciferase, using the truncated spike (S) proteins of SARS-CoV-2 [3]. The luminescence observed after treatment with luciferin reflected viral infection in cells and was highly infectious to humans (Huh7 and 293T), hamster (CHO), and monkey (Vero) cell lines [3]. Pseudotyped virus systems have been widely used for *in vitro* evaluations, such as neutralization activity [3-5]; however, whether they can also be used in animal models remains unknown.

SARS-CoV-2 uses angiotensin-converting enzyme 2 (ACE2) on the cell surface to bind and enter cells. The SARS-CoV-2 S protein does not effectively bind to mouse ACE2 [6]; therefore, sensitivity to infection is species-dependent. Thus, *in vivo* live virus infection models have been reported in SARS-CoV-2-sensitive animals such as Syrian hamsters, transgenic mice expressing human ACE2, and ferrets [7].

Animal models that can be instigated with animal BSL (ABSL)-2 will increase the opportunities for *in vivo* evaluation. Therefore, a pseudotyped virus infection was challenged in Syrian hamsters in the present study.

## Materials and Methods

### Generation of pseudotyped viruses

Pseudotyped VSVs bearing SARS-CoV-2 S proteins were generated as previously described [3]. The expression plasmid for the truncated S protein of SARS-CoV-2 and pCAG-SARS-CoV-2 S (Wuhan) was provided by Dr. Shuetsu Fukushi, National Institute of Infectious Diseases, Japan. Pseudotyped VSVs bearing envelope (G) (VSV-G) were also generated. The pseudotyped VSVs were stored at −80 °C until subsequent use.

### Hamster models

Male 6-to 10-week-old hamsters were purchased from Japan SLC Inc. (Shizuoka, Japan). All animals were housed in a pathogen-free environment at the Division of Animal Resources and Development at the University of Toyama.

After titration of the viral solution, 100 μL (7.1 × 10^5^–10^6^ RLU/hamster for SARS-CoV-2 pseudotyped virus [SARS-CoV-2pv] and 7.1 × 10^7^–10^8^ RLU/hamster for VSV-G pseudotyped virus [VSV-Gpv]) were directly inoculated into the trachea as previously described [8]. Briefly, after anesthesia with isoflurane or a mixture of medetomidine (0.15 mg/kg), midazolam (2 mg/kg), and butorphanol (2.5 mg/kg), viral solutions were administered through the trachea using an 18 G (65 mm long) catheter (TOP Co., Tokyo, Japan) with a 1 mL syringe under the assistance of an ear pick with light (Figure 1). Immediately after confirming that the solution in the syringe showed respiratory fluctuations, the viral solution was administered into the lower respiratory tract. The Ethics Review Committee for Animal Experimentation approved all experimental protocols used in the present study (Protocol Number: A2020MED-18).

**Figure 1.**
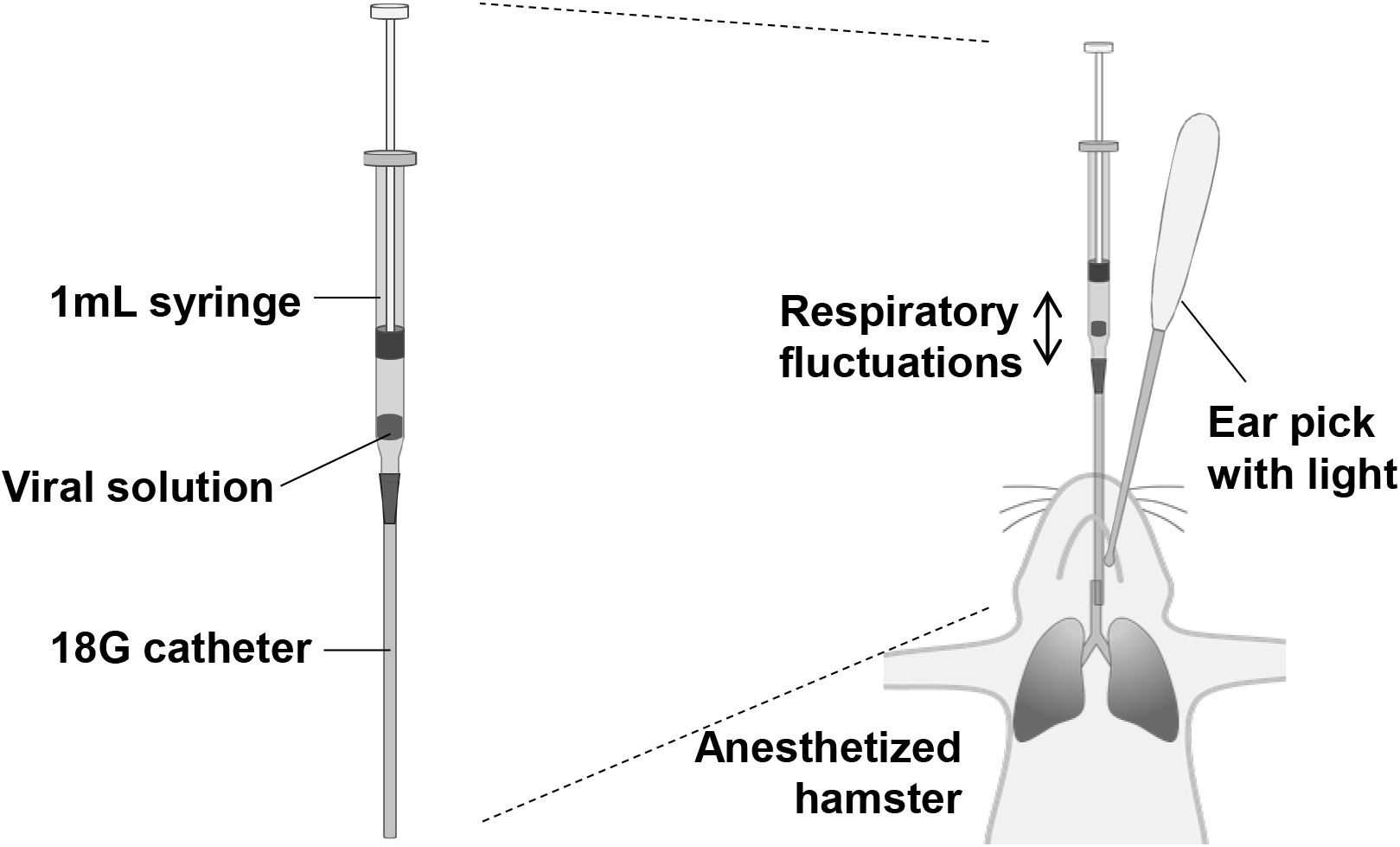
Schematic illustration of the inoculation of virus solution into the hamster airway. A viral solution (100 μL/hamster) with a small amount of air was prepared in a 1 mL syringe attached to an 18G catheter. After observing the glottis of the anesthetized hamster with an ear pick with light, the syringe set containing the viral solution was inserted through the oral cavity and advanced through the vocal cords. The respiratory fluctuations of the viral solution indicated that the tip of the catheter was in the trachea.

### Bioluminescent imaging

The hamsters were sacrificed by isoflurane and cervical dislocation 24 h after infection. The lungs were harvested and washed with phosphate-buffered saline (Nakarai Tesque, Kyoto, Japan). The lungs were incubated with 1 mg/mL of D-luciferin (Promega, Madison, WI, USA) for 5 min, and luminescence was measured using an in vivo imaging system (IVIS Lumina II, Perkin Elmer, MA, USA). Analyses were performed using Living Image 4.2 software (Caliper Life Science) to measure the light emitting from the infection sites. The luminescence from the front and back of the lungs was measured and the sum was calculated. All values are expressed as photons per second per cm^2^ per steradian (p/sec/cm^2^/sr) for each mouse.

### Quantification of VSV N expression

To evaluate viral infection, VSV N gene expression in the lungs was measured using a quantitative polymerase chain reaction [9]. The lungs were homogenized in a tube containing glass beads and 500 μL of pre-chilled phosphate-buffered saline by BeadMill 24 (Thermo Fisher Scientific, MA) at a speed of 6 m/s for 60 s. Each 20 μL of homogenized lung solution was immediately mixed with 500 μL of Isogen (Nippon Gene, Toyama, Japan) and stored at -80 °C until subsequent use. The total RNA was extracted using a QIAamp Viral RNA Mini Kit (Qiagen, Hilden, Germany) according to the manufacturer’s instructions. The total RNA was reverse transcribed into cDNA and then amplified using a Thunderbird SYBR qPCR/RT Set III (TOYOBO Co.., Tsuruga, Japan). VSV N expression was normalized to that of γ-actin [10].

### Statistical analysis

The luminescence data are expressed as the mean ± standard error of the mean (SEM). Differences between low and high SARS-CoV-2pv concentrations were examined for statistical significance using an unpaired Student’s t-test with GraphPad Prism version 8.4.3 (GraphPad Software, CA). Because high values for higher SARS-CoV-2pv concentrations were expected, a one-tailed P value of < 0.05 denoted the presence of a statistically significant difference.

## Results

### Intratracheal inoculation of SARS-CoV-2 pseudotyped virus demonstrated signs of lung infection

VSV-Gpv or SARS-CoV-2pv inoculation induced no clinical signs until 24 h after inoculation.

For VSV-Gpv, no luminescence was detected in the lungs inoculated with 7.1 × 10^6^ RLU/hamster (1-fold concentration, data not shown). Thus, higher concentrations of VSV-Gpv (10- and 100-fold concentrations) were inoculated (Figure 2A). The luminescence values of the 10- and 100-fold concentrations were 0.16 ± 0.08 × 10^7^ and 2.8 ± 0.88 × 10^7^ p/sec/cm^2^/sr, respectively (p < 0.05, n = 3 each) (Figure 2B).

**Figure 2.**
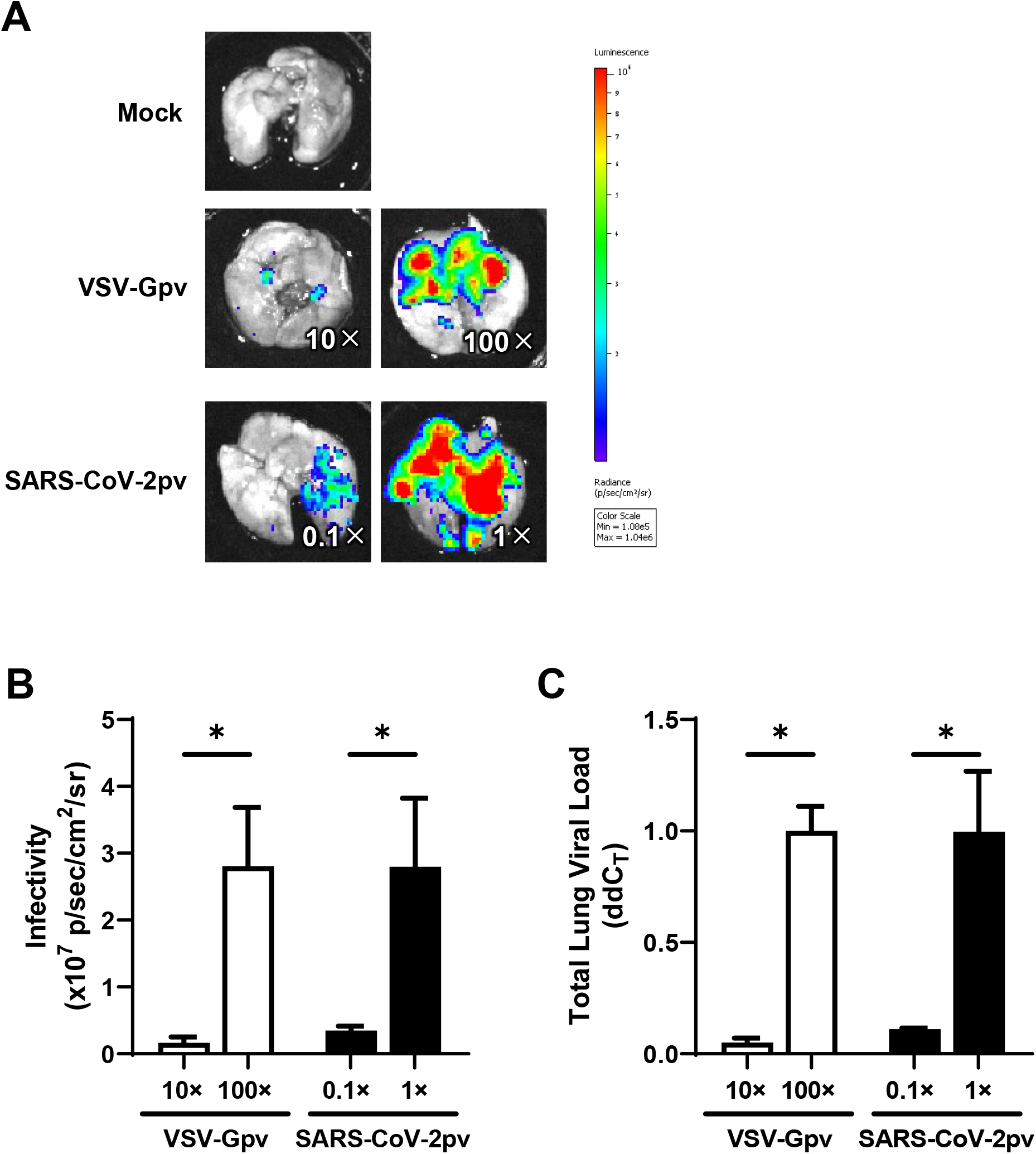
Dose-dependent infectivity after inoculation of the pseudotyped virus. **(A)** Representative lung imaging after inoculation of mock (top), VSV-Gpv (middle), and SARS-CoV-2pv (bottom). The numbers indicate the concentration ratio from the original viral solution (7.1 ×10^6^ RLU/hamster). **(B)** Infectivity after inoculation of VSV-Gpv (n = 3) and SARS-CoV-2pv (n = 3). Data are expressed as the mean of sum luminescence of the front and back. **(C)** Viral loads in lungs evaluating VSV N expression. Data are expressed as the mean of ddCT (VSV-Gpv; n = 2, SARS-CoV-2pv; n = 3). All experiments were performed 24 h after inoculation. Error bars are the SEM. * indicates p < 0.05.

In contrast, the luminescence was clearly observed when 7.1 × 10^6^ RLU SARS-CoV-2pv was inoculated (Figure 2A). The mean ± SEM of the luminescence values after inoculation of 7.1 × 10^5^ RLU/hamster (0.1-fold concentration) and 7.1 × 10^6^ RLU/hamster (1-fold concentration) were 0.35 ± 0.11 × 10^7^ and 2.80 ± 1.8 × 10^7^ p/sec/cm^2^/sr, respectively (p < 0.05, n = 3 each) (Figure 2B).

The viral loads in the lungs increased in a dose-dependent manner in VSV-Gpv and SARS-CoV-2pv inoculated hamsters (Figure 2C).

## Discussion

Our results demonstrated that the VSV-based SARS-CoV-2pv was applicable to SARS-CoV-2 respiratory infection in the hamster model. This method could be used as an alternative model for experiments in an ABSL-3 facility or to avoid handling highly infectious viruses, and it expands the possibility of analysis using equipment that cannot be used with BSL-3/ABSL-3.

A lower respiratory infection model using pseudotyped SARS-CoV-2 viruses has not been reported to date. Although a mouse model using lentivirus-based SARS-CoV-2pv following intranasal administration of human ACE2 has been reported [11], the virus did not infect the lower respiratory tract and remained around the nose. Generally, nasal inoculation is a convenient route to induce lower respiratory tract infection. However, in our preliminary experiments, sufficient luminescence was not observed with *in vivo* imaging after nasal inoculation with the pseudotyped virus (data not shown). Therefore, it was essential to approach the lower respiratory tract directly. A tracheostomy approach from the anterior neck is one possible inoculation route for pathogens into the lower respiratory tract [12]; however, our intratracheal approach was less invasive and complicated. Confirmation of respiratory fluid fluctuations provides good reproducibility for models. The time required for the inoculation was approximately 1–3 min/hamster.

Our model can be used for research on viral binding steps. For example, it would be useful for research on the treatment and prevention of viral binding and entry. However, this model is unsuitable for assessing advanced status such as severe conditions and extrapulmonary infections because of the single-round infectious virus. Because the viral solution was sprayed blindly, the infected lesion might be one-sided, and it was better to evaluate both the front and back sides. The imaging findings were consistent with VSV N expression, suggesting that *in vivo* imaging is a suitable assay for evaluating viral load. In the present study, despite being inoculated with the same infectivity VSV-Gpv *in vitro*, an approximately 100-fold concentration was required to show relative luminescence *in vivo*. This finding might be because the expression ratios of the low-density lipoprotein receptor used by VSV [13] and the ACE2 receptor used by SARS-CoV-2 differ *in vitro* and *in vivo*. However, VSV-Gpv could be used as a control in intervention studies by inoculating with a 100-fold dose, if required.

In conclusion, we successfully established a hamster model of SARS-CoV-2 respiratory infection using a VSV-based pseudotyped virus. This model can be used to evaluate the treatment and preventive potential without using highly infective pathogens and transgenic animals. Because it is not limited to ABSL-3 containment, further studies can evaluate different ideas, which will be useful for future COVID-19 studies.

## Acknowledgements

We would like to thank Makito Kaneda and Yushi Murai for their assistance with anesthesia and euthanasia. We also gratefully acknowledge Ms. Yumiko Nakagawa for secretarial assistance.

## Data availability

There is no additional data.

## Authors’ contributions

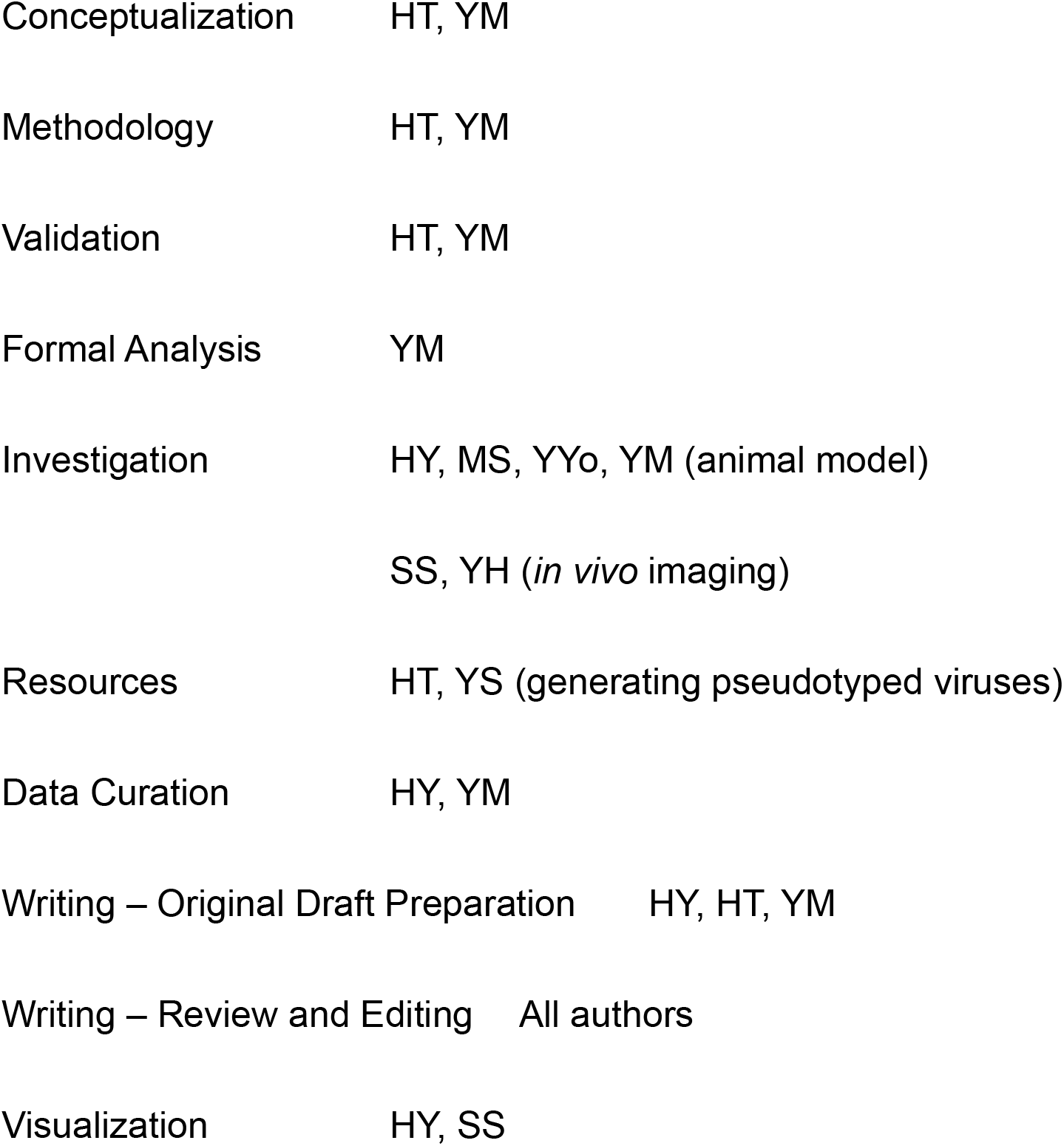
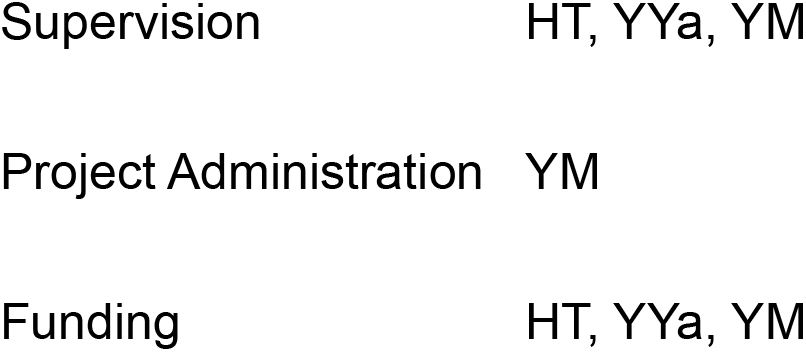

